# Probabilistic coupling of cellular and microenvironmental heterogeneity by masked self-supervised learning

**DOI:** 10.64898/2026.04.21.719876

**Authors:** Yasuhiro Kojima, Yosuke Tanaka, Haruka Hirose, Fumiko Chiwaki, Kazuya Nishimura, Shuto Hayashi, Kota Itahashi, Masato Ishikawa, Teppei Shimamura, Hiroyuki Mano

**Author notes:** Corresponding author: Yasuhiro Kojima. These authors contributed equally to this work (Yasuhiro Kojima, Yosuke Tanaka).

## Abstract

Spatial omics technologies have advanced to single-cell resolution, enabling systematic analysis of tissue microenvironments alongside cellular-state heterogeneity. However, computationally defining microenvironmental states at single-cell resolution and identifying representations most informative for biological discovery remain major challenges. Here we present Mievformer, a Transformer-based masked self-supervised framework that learns microenvironmental embeddings by encoding neighboring cellular states and relative spatial configurations to parameterize the conditional distribution of continuous cell states at central spatial positions. Through InfoNCE optimization, Mievformer learns representations that capture the relative enrichment of cell states across microenvironments, formalized as a conditional density ratio, thereby enabling probabilistic inference of the coupling between microenvironmental and cellular heterogeneity. Mievformer outperformed existing methods in niche clustering on simulated spatial transcriptomics data and achieved the highest average performance across five real datasets spanning three spatial transcriptomics platforms when evaluated using DREC, a ground-truth-free metric that most strongly correlated with ground-truth performance in simulations. Beyond conventional clustering, Mievformer enables identification of cellular subpopulations based on their microenvironmental distribution and detection of gene-expression signatures associated with colocalization of specific cell populations. Together, these results establish Mievformer as a quantitatively robust and biologically informative framework for learning microenvironment representations in spatial omics.

## 1 Introduction

Single-cell transcriptomics has revealed prevalent cellular heterogeneity across diverse biological systems, including disease states[1]. These data have enabled the reconstruction of developmental lineage hierarchies and their master regulators, as well as the identification of disease-associated cell populations that clarify cell-level pathomechanisms. However, the progression and diversification of multicellular systems, such as tumors, are driven not only by autonomous cell-state transitions but also from sequential, context-dependent cell-cell interactions that define the evolution of cellular communities[2, 3]. Thus, elucidating tissue development and disease through the framework of cellular heterogeneity requires characterising the heterogeneity of microenvironments wherein multiple cell types interact. This perspective drives omics analyses of the cell states that produce microenvironmental heterogeneity and the identification of cell populations that are enriched in specific microenvironments.

Spatial omics, conducted by measuring gene expression in its native spatial context, offers unprecedented power to dissect microenvironmental heterogeneity[4], since it enables the identification of disease-specific microenvironments [5] and improves data-driven inference of cell-cell interactions by providing spatial proximity constraints [6]. However, the notion of “microenvironmental state” remains conceptually ambiguous, hampering its computational definition and estimation. A prevalent strategy is to learn representations that directly reconstruct spot-level expression from spatial transcriptomics. For example, BayesSpace[7] assigns discrete microenvironmental states under Markov random-field constraints to enforce spatial coherence and optimise state-specific expression profiles and assignments via Markov Chain Monte Carlo. STAGATE[8] and GraphST[9] construct spatial proximity graphs and learn latent microenvironmental states using graph-based encoders. However, these approaches equate the microenvironmental state with the expression observed at each spatial location, which can be limiting for modern single-cell-resolution platforms where cells with distinctively different expression profiles such as tumor and immune cells can be adjacent to each other in a shared microenvironment. Recently, several methods have been proposed to better leverage the increasing resolution of spatial omics. NicheCompass[10] models microenvironments with a graph attention network[11] by jointly regressing the central cell expression profile and the neighbourhood-averaged expression, while regularising the latent space so that microenvironmental similarities align with the spatial proximity graph. Banksy[12] derives microenvironmental representations by augmenting neighbourhood-averaged expression with spatial expression gradients, thereby capturing features that mean profiles alone miss. CellCharter[13] constructs a spatial neighbor graph via Delaunay triangulation and aggregates neighborhood features across multiple graph layers, enabling the identification of spatial cell niches at multiple scales. Nevertheless, because these approaches rely on expression aggregated over user-defined neighbourhoods, their outputs are sensitive to the neighbourhood definition and, in practice, attenuate the single-cell resolution of the measurements.

Here, we present Mievformer, a Transformer-based masked self-supervised framework that learns microenvironmental embeddings by encoding neighboring cellular states and relative spatial configurations. Rather than regressing central expression profiles or neighborhood-aggregated features, Mievformer parameterizes the conditional distribution of continuous cell states at central spatial positions. Through InfoNCE [14] optimization, it learns representations that capture the relative enrichment of cell states across microenvironments, formalized as a conditional density ratio, thereby enabling probabilistic inference of the coupling between microenvironmental and cellular heterogeneity. This framework supports analyses beyond conventional clustering, including single-cell-resolution characterization of microenvironmental preferences and identification of gene-expression signatures associated with colocalization of specific cell populations. Together, these capabilities establish Mievformer as a quantitatively robust and biologically informative framework for spatial omics analysis.

## 2 Results

### 2.1 Overview of Mievformer

Mievformer learns microenvironmental representations by encoding the cellular states and spatial configurations of neighboring cells using a Transformer-based architecture with masked self-supervised learning. Specifically, it masks the cell state at a central cell position and employs InfoNCE loss to maximize the mutual information between the inferred microenvironmental embedding and the observed central cell state. This contrastive learning objective yields embeddings that parameterize the conditional density ratio of cell states in local tissue contexts (see the Methods section for details).

Formally, given spatial transcriptomics data comprising gene expression *y*, low-dimensional expression embeddings *z* obtained via Principal component analysis (PCA), and spatial coordinates *x*, Mievformer operates as follows: For each cell *i* with spatial neighborhood *N* (*i*), the microenvironmental embedding is computed as:

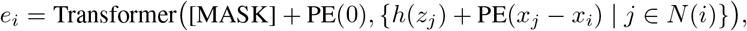

where *h* is a learned projection of cell embeddings and PE(*·*) denotes the sinusoidal positional encoding of relative spatial coordinates. The model defines a score function *s*(*e*_*i*_, *z*_*j*_) and computes the probability *P*_*ij*_ that the cell state *z*_*j*_ would be observed at the masked position via softmax normalization across all cells in the mini-batch. Training maximizes *P*_*ii*_, the likelihood of the actual observed cell state *z*_*i*_, which is equivalent to minimizing the InfoNCE loss. This optimization learns embeddings such that the score function approximates the conditional density ratio *p*(*z*|*e*)*/p*(*z*). Once trained, Mievformer can compute microenvironmental embeddings for any spatial position and derive the stochastic association between cell states and microenvironments from the learned density ratio.

Beyond conventional clustering-based characterization of microenvironmental heterogeneity, the density ratio learned by Mievformer enables several downstream analyses through probabilistic inference. First, by applying density-ratio-based Bayesian inversion to the learned ratio *p*(*z*|*e*)*/p*(*z*), *p*(*e*|*z*)*/p*(*e*) is obtained, which represents the posterior density ratio of microenvironments given a cell state. Aggregating these ratios over microenvironmental clusters yields single-cell-resolution microenvironmental membership profiles, which can be used to identify cell subpopulations with specific microenvironmental preferences. Second, calculating the expectation of the density ratio *p*(*z*|*e*)*/p*(*z*) across all cell states belonging to a specific cell population by Monte-Carlo approximation, the density of that population in microenvironment *e* is obtained. Correlation analysis with this density reveals the genes up- or down-regulated given the colocalization with the target cell popution. Together, these capabilities establish Mievformer as a quantitative framework for clarifying the joint structure of cellular and microenvironmental heterogeneity in spatial omics data (Figure 1).

**Figure 1:**
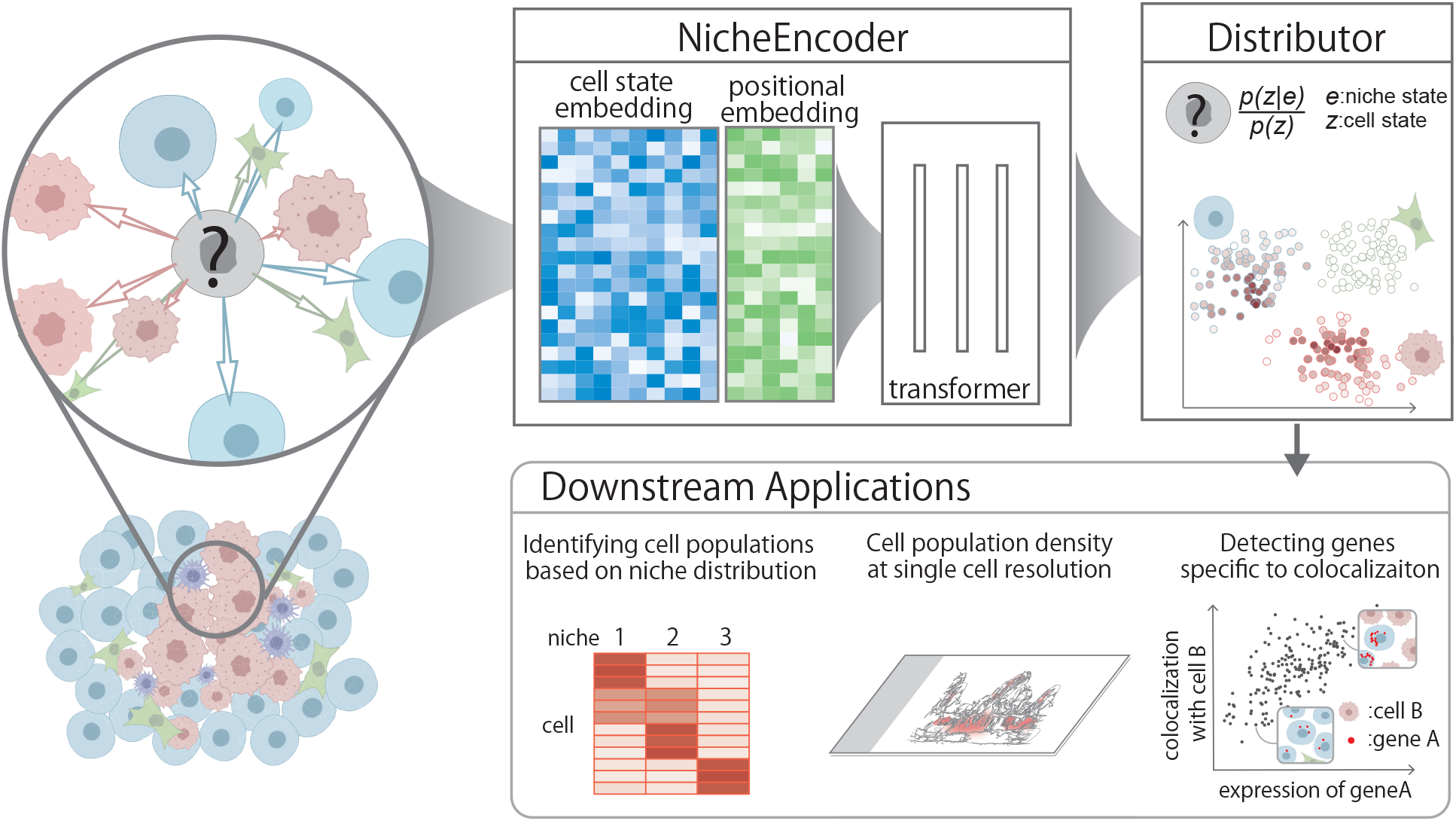
Overview of the Mievformer framework. **(Left)** Schematic of spatial tissue context: neighboring cells surrounding a central cell (marked with “?”) whose cell state is masked and microenvironmental state is to be inferred. **(Center)** The NicheEncoder module receives cell state embeddings and positional embeddings of neighboring cells and processes them through a Transformer architecture to produce a microenvironmental embedding. **(Right)** The Distributor module computes the conditional density ratio *p*(*z*|*e*)*/p*(*z*), quantifying how much more likely each cell state is to be observed in the inferred microenvironment relative to the marginal distribution. **(Bottom)** Downstream applications enabled by the learned density ratios: identifying cell subpopulations based on their microenvironmental distribution profiles, estimating cell population density at single-cell resolution, and detecting genes specific to colocalization patterns.

### 2.2 Performance superiority in deriving representations of microenvironment in simulated data

To quantitatively assess the performance of Mievformer, we applied it to simulation data configured with multiple patterns, each comprising three ground-truth microenvironmental regions (inner, boundary, and outer), and compared its performance against existing methods (Figure 2). The simulation consisted of multiple concentric annular patterns, each composed of these three region types and assigned a distinct, pattern-specific cell-cluster composition. These repetitive lumen-like patterns were designed to mimic tissue architectures observed in biological structures such as thyroid follicles, renal tubules, and seminiferous tubules. For each spatial spot, a cluster label was first determined according to the composition of its assigned region (inner, boundary, or outer), and then an expression profile was randomly sampled from cells belonging to that cluster.

**Figure 2:**
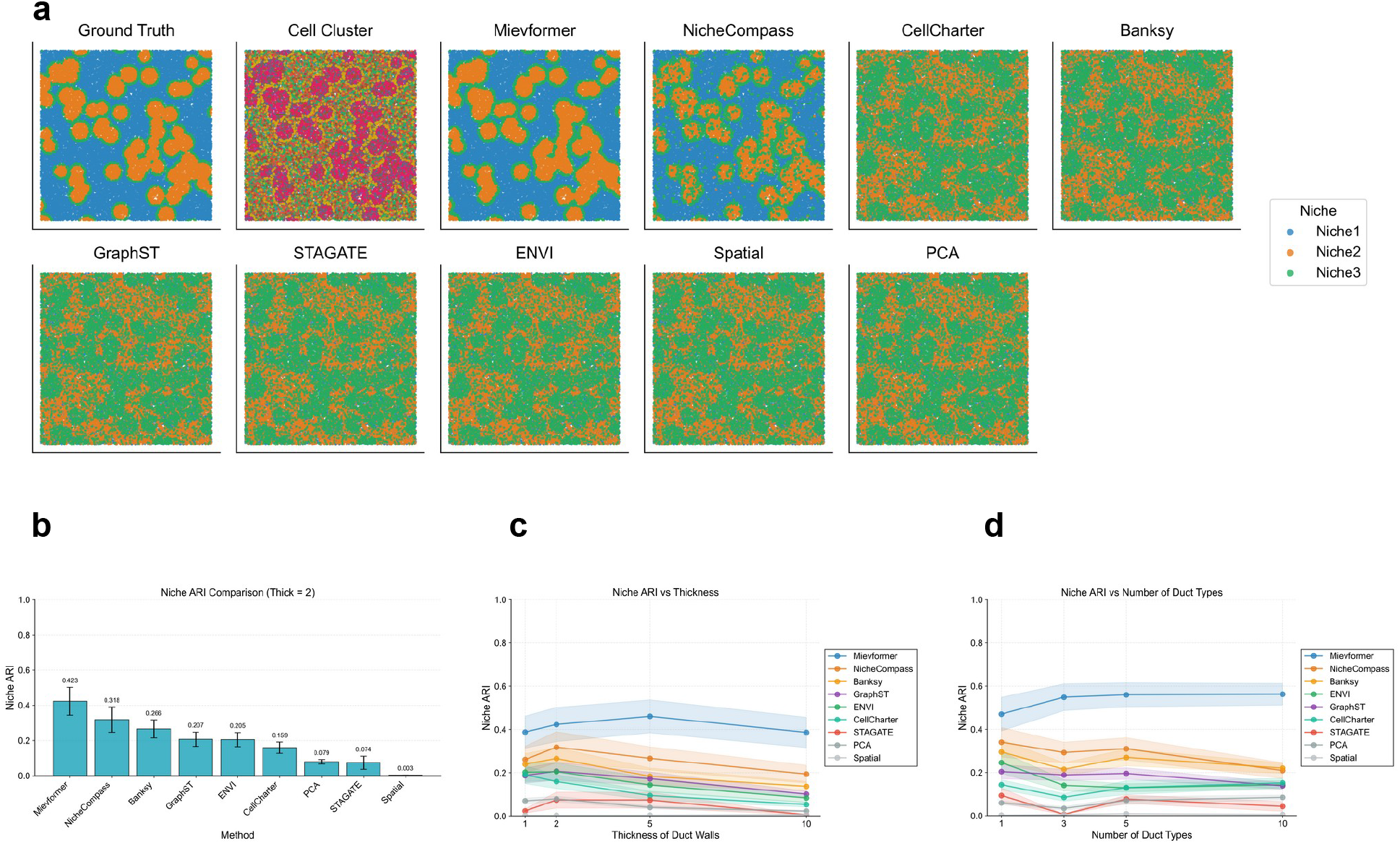
Simulation Comparisons for Niche Clustering Methods. **(a)** Spatial niche pattern comparison using Hungarian algorithm for optimal cluster matching across methods (Mievformer, NicheCompass, CellCharter, Banksy, GraphST, STAGATE, ENVI, Spatial, PCA). Ground truth niche patterns and cell clusters are also presented as references. **(b)** Adjusted Rand Index (ARI) comparison across clustering methods at standard duct wall thickness (thick=2) with standard errors. **(c)** Niche ARI performance as a function of duct wall thickness. The shades represent the standard errors. **(d)** Niche ARI performance as a function of number of duct types (categorical complexity). The shades represent the standard errors.

We applied Mievformer, NicheCompass, CellCharter, Banksy, COVET, STAGATE, and GraphST to the simulated spatial transcriptomics data. The microenvironmental representations obtained from each method were partitioned into three clusters using *k*-means clustering. As baselines, we also performed clustering on PCA expression profiles and spatial coordinates. Qualitative comparison of the spatial distributions of estimated microenvironmental clusters with ground-truth regions revealed that Mievformer most faithfully reproduced the three-layered structure of inner, boundary, and outer regions (Figure 2a). Quantitative evaluation using the Adjusted Rand Index (ARI) confirmed that Mievformer achieved superior agreement with ground truth compared to that of all competing methods (Figure 2b). All methods outperformed the PCA and spatial coordinate baselines. To assess robustness, we further evaluated performance across varying simulation parameters, including boundary layer thickness and number of distinct cell composition patterns. Although ARI values were reduced for all methods under conditions with fewer patterns or thinner boundary layers, Mievformer consistently achieved the highest performance across all tested conditions (Figure 2c,d).

### 2.3 Benchmarking microenvironmental representations on real datasets

Next, we compared the performance of microenvironmental representation methods on five real spatial transcriptomics datasets: Xenium datasets from pancreatic, lung, and glioblastoma cancers, a Stereo-seq mouse brain dataset[15], and a Visium HD mouse brain dataset. We applied *k*-means clustering to the microenvironmental representations obtained by each method and qualitatively compared their spatial distributions. Relative to clusters derived from PCA coordinates, all methods exhibited more spatially coherent patterns; however, qualitative assessment alone was insufficient to determine superiority among methods (Figure 3a).

**Figure 3:**
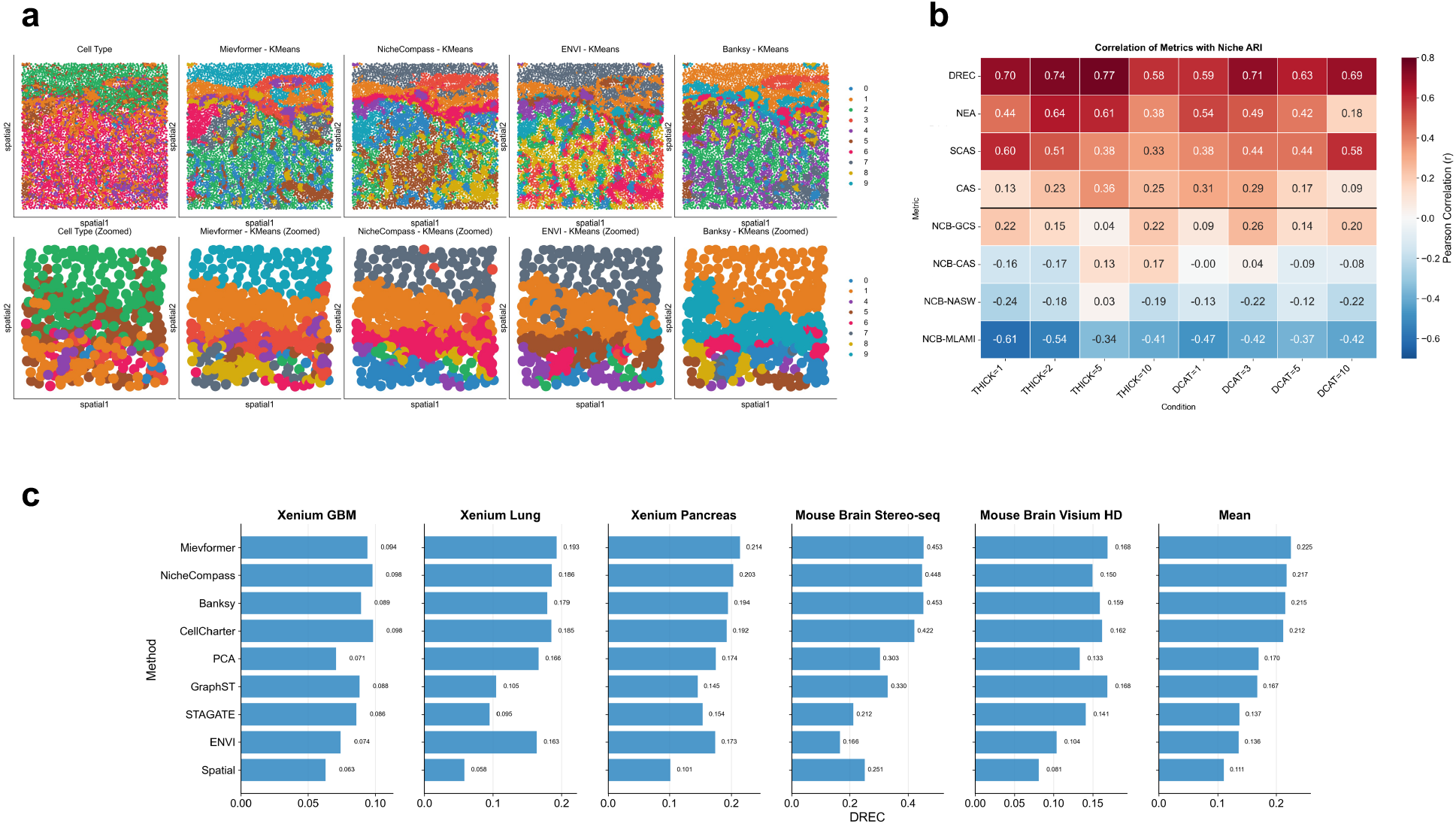
Real Data Comparisons Across Tissue Types. **(a)** Spatial visualization comparing k-means clustering of niche representations from methods (Mievformer, NicheCompass, CellCharter, Banksy, GraphST, STAGATE, ENVI, Spatial, PCA). cell type annotations (left). **(b)** Heatmap showing the Pearson correlation between eight evaluation metrics (CAS, SCAS, NEA, DREC, NCB-GCS, NCB-CAS, NCB-NASW, NCB-MLAMI) and ground-truth Niche ARI across varying simulation parameters. **(c)** Horizontal bar graphs showing dual recovery (DREC) metrics across methods and five datasets: Xenium GBM, Xenium Lung, Xenium Pancreas, Mouse Brain Stereo-seq, and Mouse Brain Visium HD.

Although multiple quantitative metrics have been proposed and employed in prior studies of microenvironmental representation, their validity has not been rigorously established. To address this, we evaluated the correlation between several metrics and ground-truth performance (Niche ARI) in simulation data. Specifically, we examined one existing metric (CAS), three novel metrics we propose (SCAS, NEA, and DREC), and four metrics from the NicheCompass benchmarking module[10] (NCB-GCS, NCB-CAS, NCB-NASW, NCB-MLAMI). Our implemented metrics are summarized as follows:

#### CAS (Cell Type Affinity Similarity)

Measures the correlation between cell-type co-localization patterns in microenvironmental embedding neighborhoods and in spatial neighborhoods, thereby quantifying the preservation of spatial co-localization. However, this metric can assign high scores to representations that closely approximate spatial coordinates themselves.

#### SCAS (Shuffled Cell Type Affinity Similarity)

After clustering microenvironmental states, spatial coordinates are randomly permuted within each cluster, and the correlation of cell-type co-localization patterns with the original coordinates is computed. This metric tests whether co-localization patterns are retained under shuffling, mitigating the bias favoring spatial-coordinate-based embeddings.

#### NEA (Neighbourhood Enrichment Accuracy)

Evaluates consistency between cell-type composition ratios in microenvironmental clusters and spatial co-localization. Defined as the Pearson correlation between an enrichment matrix *E*_sp_ computed from spatial nearest neighbours (2-nearest neighbours) and an expected enrichment matrix *E*_niche_ = *P · P*^*T*^ derived from the cell-type composition matrix *P* within microenvironmental clusters.

#### DREC (Dual Recovery)

Quantifies the proportion of expression-similar cells whose nearby spatial neighbors belong to the same microenvironmental cluster as the original cell. Specifically, for each cell, we select the *k*-th nearest cell in expression space (based on the top 20 principal components, with *k* = 10) and evaluate whether the *k*-nearest spatial neighbor (*k* = 10) of that selected cell belongs to the same microenvironmental cluster as the original cell.

The comparison of these metrics with ground-truth Niche ARI in simulation data revealed that DREC exhibited the strongest correlation across varying simulation parameters, including duct thickness and number of duct categories (Figure 3b). Conversely, CAS showed the second-lowest correlation. Notably, the baseline representation using spatial coordinates alone achieved the highest CAS value, confirming our concern that CAS favors proximity-based features. Although SCAS exhibited moderate correlation, performance of multiple methods converged near ceiling values, precluding Mievformer’s superiority observed in ARI and qualitative analyses. Both SCAS and NEA, similar to CAS, evaluate coarse-grained inter-cell-type co-localization rather than single-cell-resolution microenvironmental consistency, which likely explains DREC’s superior discriminative power. The four NCB metrics generally exhibited weak or negative correlations with Niche ARI, with NCB-MLAMI showing a strongly negative correlation (−0.34 to −0.61). Based on these findings, we adopted DREC as the primary performance metric for subsequent real-data analyses.

Finally, we benchmarked all methods on five real datasets using DREC (Figure 3c). Mievformer achieved the highest DREC score in four of the five datasets, with only the Xenium GBM dataset being an exception. In the Visium HD Mouse Brain dataset, Mievformer tied with GraphST for the top position. Across all five datasets, Mievformer attained the highest mean DREC score. Notably, the PCA baseline, which is simple expression-based clustering, also exhibited high DREC scores, with only Mievformer, NicheCompass, Banksy, and CellCharter exceeding it. This likely reflects the biological tendency for expression-similar cells to co-localize spatially. Nevertheless, these results demonstrate that Mievformer captures microenvironmental structure more faithfully than existing methods, including expression-based baselines, suggesting its representations better reflect latent microenvironmental organization.

### 2.4 Elucidating microenvironmental diversity in lung cancer

To qualitatively assess whether Mievformer’s microenvironmental embeddings capture biologically meaningful microenvironmental diversity, we performed detailed interpretation of Mievformer outputs on lung cancer Xenium data (Figure 4). Clustering of the microenvironmental embeddings identified twelve clusters with specific cell-type compositions (Figure 4a,b). Five clusters (Niche 4, 5, 6, 7, and 10) were dominated by tumor cells, whereas Niche 1 predominantly was occupied by monocytes/macrophages, Niche 0 was T cell-enriched, and Niches 2, 3, and 11 were fibroblast-dominant niches. Notably, even among the tumor-dominant clusters, Mievformer resolved fine-grained microenvironmental differences: Niche 4 comprised >80% tumor cells, Niche 5 contained 20% fibroblasts alongside tumor cells, and Niche 7 resembled Niche 4 but included 10% endothelial cells.

**Figure 4:**
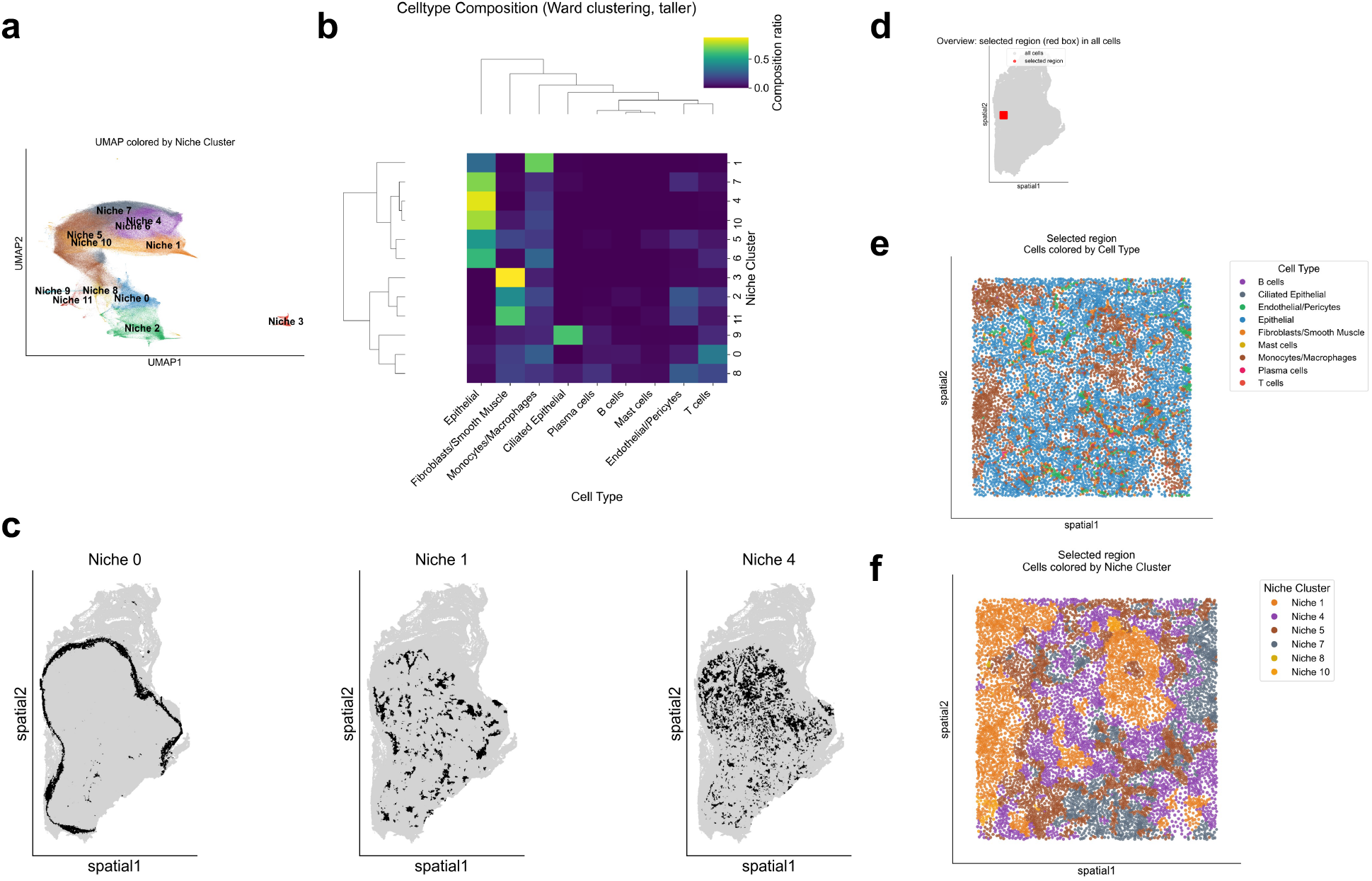
Niche Heterogeneity Analysis in Lung Tissue. **(a)** UMAP visualization of Mievformer’s latent representation for Xenium lung cells, colored by niche cluster assignment. **(b)** Hierarchical clustering heatmap showing cell type composition across niche clusters. **(c)** Spatial plots for niche clusters 0, 1, and 4. Black points indicate cells belonging to the target niche cluster; light gray points show all other cells. **(d)** Overview of lung tissue section with selected region marked by black rectangle. **(e)** Cell type spatial plot of the selected region. **(f)** Niche cluster spatial plot of the selected region.

Spatial analysis revealed that tumor-dominant niche clusters (4, 5, 6, 7, and 10) were localized to the tumor core, fibroblast-enriched clusters (2 and 3) occupied stromal regions, and immune-enriched clusters were concentrated at tumor-stroma boundaries (Figure 4d). Interestingly, macrophage-dominated Niche 1 was distributed throughout the tumor interior. Direct comparison of the spatial patterns of Niches 4 (tumor-enriched) and 1 (macrophage-enriched) confirmed that Mievformer successfully distinguished these spatially intermingled microenvironments (Figure 4c).

Together, these results demonstrate that Mievformer embeddings enable biologically interpretable dissection of microenvironmental heterogeneity.

### 2.5 Identification of cell subpopulations based on microenvironmental distribution

Mievformer enables the characterization of cell-type subpopulations based on their microenvironmental distributions by computing membership probabilities to microenvironmental clusters for each individual cell. To quantitatively validate the membership probabilities assigned by Mievformer, we aggregated the estimated probabilities for each combination of microenvironmental clusters and cell types (Figure 5b). For all the cell types, the membership probability to the corresponding niche cluster was consistently the highest among microenvironmental clusters containing *≥* 5% of cells from the given cell type (Figure 5b). While fibroblasts exhibited highly specific cluster assignments, we observed the some combinations of microenvironmental clusters and cell types exhibited relatively low specificity to corresponding microenvironmental clusters. For instance, tumor cells in Niche 5 showed substantial assignment not only to Niche 5 but also to Niche 7, and macrophages in Niche 10 similarly exhibited increased assignment to Niche 7.

**Figure 5:**
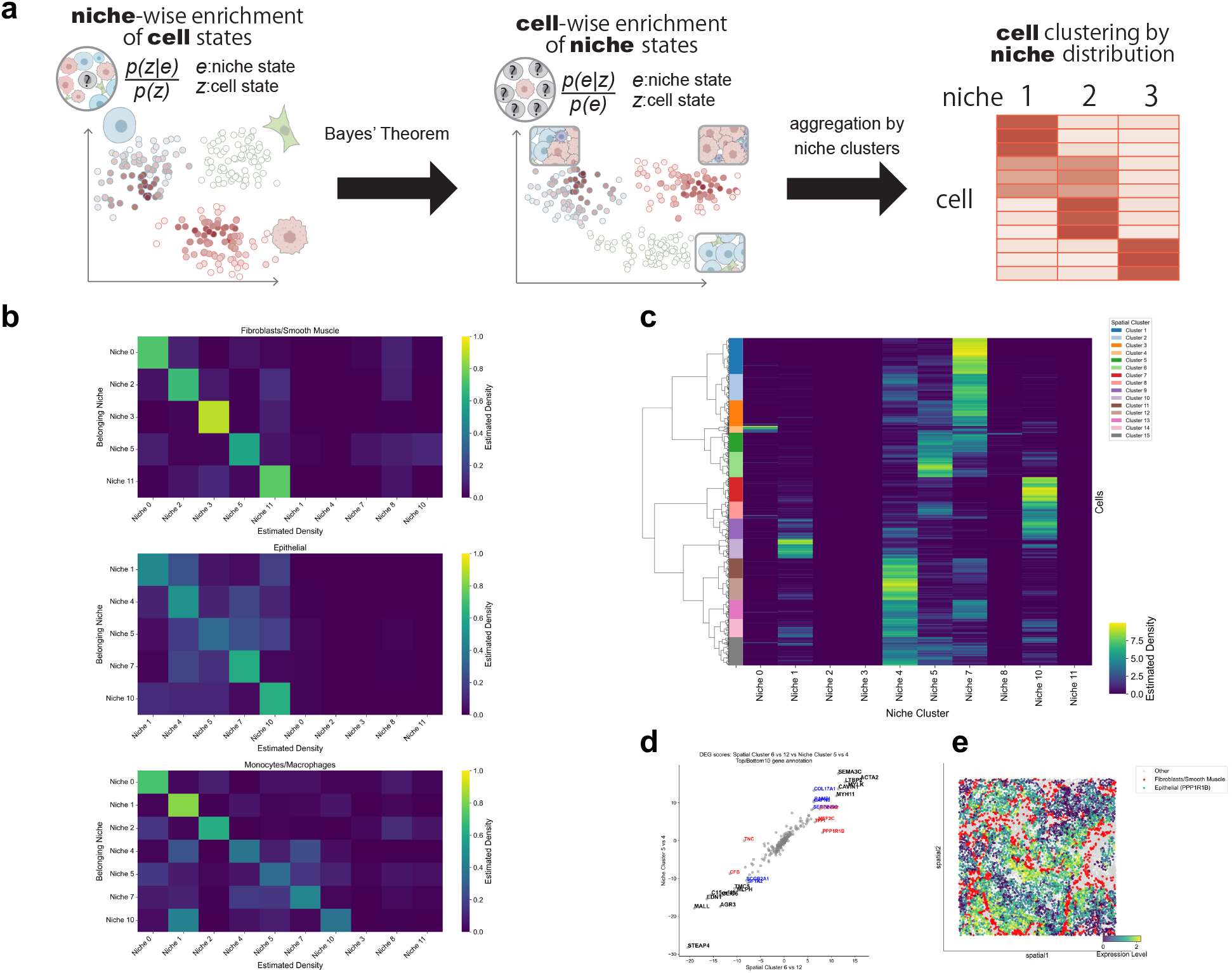
Cell-wise posterior distribution across microenvironments. **(a)** Schematic overview of spatial-cluster-based cell subpopulation identification. For each cell, membership probabilities across microenvironmental clusters are computed by Mievformer, and hierarchical clustering on these distribution profiles defines microenvironment-specific subpopulations (spatial clusters; labeled Dist N). **(b)** Heatmap showing niche-cluster-wise aggregated posterior distribution across nine niche clusters for Fibroblasts/Smooth Muscle, Epithelial, and Monocytes/Macrophages. **(c)** Single-cell-wise posterior distribution across microenvironments for epithelial cells. Rows are ordered by hierarchical clustering; row colors indicate the spatial cluster assignment derived from this clustering. **(d)** Scatter plot of differentially expressed gene scores. X-axis: DEG scores between spatial clusters (Dist 6 vs 12), Y-axis: DEG scores between niche clusters (Niche 5 vs 4). DEG scores were calculated using the Wilcoxon rank-sum test, implemented in Scanpy’s `rank_genes_groups` function. **(e)** Spatial expression plot of PPP1R1B gene. Red marks location of fibroblasts.

To investigate whether these differences in mean assignment probability reflected overall differences in cluster specificity or intra-cluster variation, we examined the distribution of membership probabilities within each microenvironmental cluster for each cell type. Clusters with specific assignments displayed sharp peaks near probability 1.0, whereas clusters with non-specific assignments, although containing cells with high assignment probabilities to the corresponding cluster, exhibited broad distributions with substantial variance (Supplementary Figure 2). These results indicate that Mievformer’s membership probabilities to microenvironmental clusters correctly capture the relationship between expression variation within cell types and distribution to microenvironmental clusters, suggesting that for certain cell types and microenvironmental clusters, there exist cells specifically enriched in particular microenvironments and those that are not. To leverage this distribution across microenvironmental clusters as a novel cellular feature encoding niche specificity, we performed hierarchical clustering based on microenvironmental membership profiles of tumor cells (Figure 5c). This approach stratified subpopulations (Dist 1, …, and Dist 15) by microenvironmental specificity. For instance, among cells with high assignment to Niche 5, we identified Dist 5, which also showed increased assignment to Niche 7, and Dist 6, which exhibited specific assignment exclusively to Niche 5. Similarly, we identified tumor cell spatial clusters Dist 7, 10, 4, 1, and 6, each specifically enriched in Niche 10, 1, 0, 7, and 5, respectively, where tumor cells are moderately abundant. To assess the biological utility of this microenvironment-specific cell population identification, we performed comparative differential expression gene (DEG) analyses. We contrasted two approaches: (1) comparing tumor cells in each niche (Niches 10, 1, 0, 7, and 5) versus those in Niche 4 (the cluster with the highest tumor cell proportion), and (2) comparing tumor cells in the corresponding spatial clusters (Dist 7, 10, 4, 1, and 6) versus those in Dist 12 (which specifically localizes to Niche 4) (Figure 5d, Supplementary Figure 4).

While the two approaches yielded broadly similar trends across most niches, spatial clustering revealed distinct expression signatures not detected by niche-based analysis. For example, PPR1B, which was not identified as DEGs between Niche 5 and 4, showed markedly differential expression between their corresponding spatial clusters Dist 6 and 12. Given that Niche 5 is characterized by abundant fibroblasts alongside tumor cells and macrophages, we examined the spatial expression patterns of PPR1B in tumor cells relative to fibroblast distribution (Figure 5e). PPR1B exhibited markedly increased expression in tumor cells surrounding fibroblasts. These results demonstrate that extracting microenvironment-specific cell populations using Mievformer’s membership probabilities enables more sensitive detection of expression changes associated with microenvironmental differences than that of comparative analysis based solely on microenvironmental clusters.

### 2.6 Single-cell resolution analysis of microenvironmental colocalization

Mievformer enables the estimation of the conditional probability that any cell state would be observed in each microenvironment. This capability allows quantitative assessment of whether cells from specific populations of interest could be distributed to particular spatial locations. By computing the probability of target cell population existence at each single-cell location, we can quantify the likelihood of colocalization with the target population, which is the probability of residing in the same microenvironment. Using these single-cell colocalization scores, we can identify genes whose expression is up-regulated given the colocalization with target populations.

We applied this framework to Xenium data of pancreatic cancer, computing microenvironmental states and cell-type-specific existence probabilities (Figure 6b–d). The estimated existence probability for each cell type was consistent with its observed spatial distribution. Quantitatively, for all cell types, the averages of the estimated density in microenvironments where that cell type was present exceeded twice the density averaged across all microenvironments (Figure 6e). These results demonstrate that Mievformer’s density estimates accurately reflect the spatial distribution of actual cell populations. Examining the histograms of cell-type densities (Figure 6f) revealed distinct colocalization behaviors: while microenvironments with estimated cancer cell density exceeding 0.99 represented a substantial fraction of cancer-cell-containing microenvironments, such highly exclusive microenvironments were nearly absent for pancreatic stellate cells. This indicates that while cancer cells frequently occupy isolated microenvironments, pancreatic stellate cells almost invariably colocalize with other cell types at close range.

**Figure 6:**
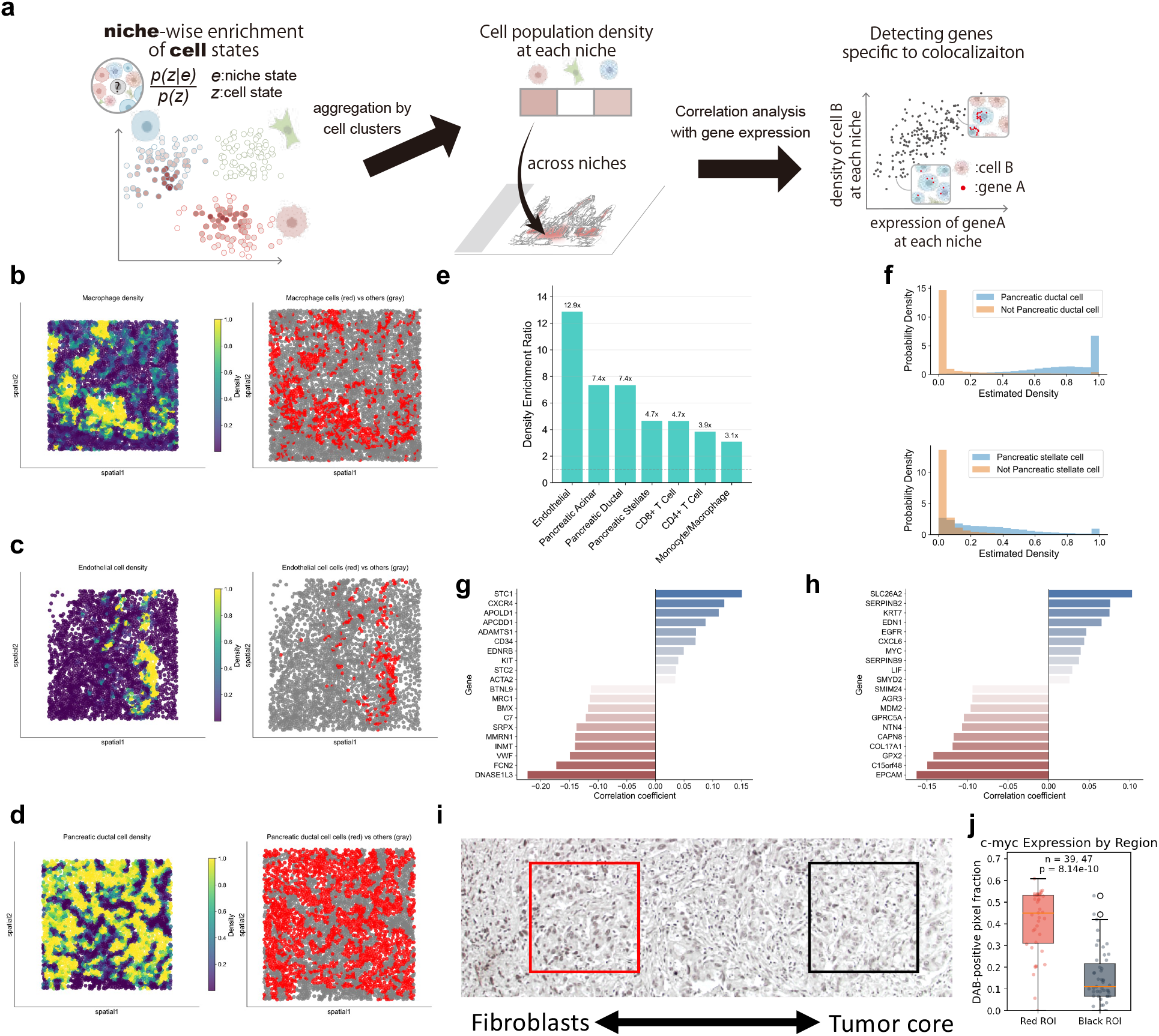
Colocalization analysis by population density estimation in pancreatic cancer. **(a)** Schematic overview of single-cell colocalization analysis. At each single-cell location, Mievformer estimates the conditional existence probability of each target cell population, from which density-based colocalization scores are computed and correlated with gene expression to identify colocalization-associated genes. **(b–d)** Spatial distribution of cell types in pancreatic cancer tissue. Left: estimated population density for Mievformer’s latent representation at each single cell spot. Right: cell positions (red=target cell type, gray=others). **(b)** macrophages, **(c)** endothelial cells, **(d)** pancreatic ductal cells. **(e)** Cell-type density enrichment barplot. For each target cell population, the bar shows the fold change of the estimated population density within that population relative to the mean estimated population density across other cells. **(f)** Histograms of estimated population density for pancreatic ductal cells (top) and pancreatic stellate cells (bottom), comparing the target cell population against all other cells. **(g)** Correlation between pancreatic ductal cell density and gene expression of endothelial cell. Only top and bottom 10 genes are shown. **(h)** Correlation between stellate cell density and gene expression of pancreatic ductal cell. Only top and bottom 10 genes are shown. **(i)** Representative immunohistochemistry image of a PDAC sample stained with c-Myc antibody, showing the locations of the analyzed regions of interest (ROIs). Brown staining indicates c-Myc positive cells. **(j)** DAB-positive pixel fraction per nucleus in fibroblast-proximal (red, *n* = 39) and tumor core (black, *n* = 47) regions. Box plots show median and interquartile range. Mann–Whitney *U* test, *p* = 8.14 × 10^−10^.

Finally, we used estimated cell densities to identify colocalization signatures with specific cell populations. For endothelial cells, we computed correlations between tumor cell density and gene expression (Figure 6g). Genes exhibiting the top10 positive correlations included CXCR4, which is specifically expressed in tip cells extending into tumor regions during angiogenesis[16], and APOLD1, which contributes to the angiogenic process[17]. These results suggest the successful capture of the expression signature of tip cells expected in peritumoral angiogenesis. For the analysis of colocalization between tumor cells and pancreatic stellate cells, we analyzed correlations between stellate cell density and gene expression in tumor cells (Figure 6h). Among genes exhibiting the top10 positive correlations, MYC has been reported to be associated with EMT in pancreatic cancer[18]. Furthermore, we validated this finding by quantifying c-Myc expression in fibroblast-proximal and tumor-core regions of PDAC sections using immunohistochemistry (Figure 6i,j). Nuclear-level quantification revealed significantly higher c-Myc expression in the fibroblast-proximal region compared with the tumor core (median DAB-positive fraction 0.45 vs. 0.11; Mann– Whitney *U* test, *p* = 8.14 × 10^−10^), consistent with a role for fibroblast-derived signals in modulating tumor cell proliferative capacity[19]. Together, these results demonstrate that Mievformer’s density-based framework enables the systematic characterization of how colocalization with specific cell populations influences expression profiles.

## 3 Discussion

In this study, we present Mievformer, a novel framework for estimating microenvironmental states as parameterizations of the conditional distribution of continuous cell states, and demonstrate its utility through simulation and real-data analyses. First, we quantitatively evaluated Mievformer’s performance relative to existing methods using simulated and real datasets. In simulations with known ground truth, Mievformer consistently achieved superior performance across a wide range of parameter settings. When applied to five real spatial transcriptomics datasets across Xenium, Stereo-seq, and Visium HD platforms, Mievformer achieved the highest average performance on our proposed evaluation metric, ranking first in four of five datasets. Beyond performance benchmarking, we demonstrated that the probabilistic coupling between cell and microenvironmental states learned by Mievformer enables biologically meaningful discovery and hypothesis generation. We established two downstream analytical frameworks: (1) cell clustering based on microenvironmental distribution profiles, and (2) identification of colocalization signatures through correlation with cell-population density estimates. Both approaches successfully recapitulated established knowledge of tumor microenvironments, validating their effectiveness for dissecting microenvironmental cell communities from single-cell spatial transcriptome data.

Since the emergence of spatial transcriptomics, numerous methodologies have characterized microenvironmental states from spatial omics data. Early spatial transcriptomics platforms lacked single-cell resolution, prompting methods that primarily sought faithful representations of spot-level expression profiles while incorporating spatial constraints to enforce consistency across neighboring spots[20]. However, this paradigm becomes problematic at single-cell resolution, where microenvironments often comprise local mixtures of transcriptionally distinct cell types, requiring representations that capture context beyond individual expression profiles. To address these limitations, recent approaches have aimed to derive low-dimensional representations from spatially aggregated features, such as neighborhood-averaged expression[10, 12]. While this strategy enables spatially coherent microenvironmental estimation even at single-cell resolution, it effectively reduces the resolution of the data to a user-defined neighborhood scale, discarding the fine-grained spatial information already present in the measurements. Conversely, Mievformer learns microenvironmental representations as parameterizations of the conditional distribution of continuous cell states at focal positions. Although computed from spatial neighborhoods, Mievformer explicitly incorporates relative spatial coordinates within those neighborhoods, enabling microenvironmental state estimation at a granularity finer than that imposed by fixed neighborhood sizes. This approach fully leverages the high spatial resolution of modern platforms and is expected to underlie Mievformer’s superior performance in microenvironmental characterization relative to existing methods.

Self-supervised learning approaches for microenvironmental state estimation have been previously proposed [10]; however, these methods fundamentally differ from Mievformer in their learning objectives and resulting representations. Some approaches reduce to regression of central cell expression profiles[21], while others frame the problem as probabilistic deconvolution of cell type proportions at each spatial location[22]. In contrast, Mievformer parameterizes the conditional probability distribution of continuous cell states at the central position, enabling probabilistic inference of the coupling between microenvironmental and cellular heterogeneity at single cell and single spot resolution. This probabilistic framework yields bidirectional inference capabilities: given a microenvironmental state, Mievformer estimates the distribution of cell states at the corresponding microenvironment; conversely, given a cell state, it computes the posterior distribution over microenvironments. These capabilities enable downstream analyses that are not readily accessible with existing methods, including estimating cell-population densities in arbitrary microenvironments and single-cell-resolution clustering based on microenvironmental membership profiles. Herein, we demonstrated the biological utility of these analyses through multiple case studies. In lung cancer data, clustering analysis based on niche distribution revealed increased PPR1B expression in tumor cells residing in fibroblast-enriched microenvironments. In pancreatic cancer data, correlation analysis between endothelial cell expression and tumor cell density identified CXCR4 upregulation in endothelial cells colocalizing with tumors, recapitulating the known expression signature of angiogenic tip cells. Together, these results establish that Mievformer’s probabilistic coupling of cell states and microenvironments provides a distinct utility for dissecting cellular heterogeneity which correlated with microenvironmental context.

Although the estimation of microenvironmental states offers significant biological utility, quantitative performance evaluation remains challenging, similar to unsupervised representation learning in other domains. NicheCompass addressed this challenge by aggregating multiple evaluation metrics to ensure robust performance assessment[10]. However, the validation of individual metrics primarily relied on their theoretical foundations rather than empirical verification. In this study, we explicitly assessed the validity of the metrics using simulation data with known ground truth. We found that some of the metrics assigned higher scores to naive baselines such as spatial coordinates or raw expression profiles instead of the methods specialized for microenvironmental representation learning, including Mievformer. Notably, these same metrics exhibited weak correlations with ground-truth performance measures. In contrast, this study introduced DREC, which demonstrated consistently strong correlation with ground-truth metrics across diverse simulation conditions. We attribute DREC’s superior discriminative power to its ability to evaluate microenvironmental consistency at single-cell resolution by directly operating on cell embeddings, thereby avoiding the information loss inherent in cluster-aggregated evaluations of cell-type composition. This finer-grained approach enables DREC to more sensitively capture microenvironmental structure than metrics that assess only coarse cell-type colocalization patterns.

Although Mievformer demonstrates superior performance in single-cell-resolution microenvironmental state estimation, certain limitations must be acknowledged. First, the current framework is optimized for characterizing spatially contiguous microenvironments and may not fully capture higher-order community structures, such as hierarchically nested or spatially interleaved microenvironmental patterns. For instance, CellCharter[13] captures multi-scale neighborhood structure through hierarchical graph-layer aggregation. Extending Mievformer to model such hierarchical organization would require architectural modifications to explicitly represent multi-scale spatial dependencies. Another important consideration is the extension to multi-sample spatial transcriptomics analysis. Mievformer is primarily designed to characterize microenvironmental heterogeneity within individual tissue sections, but recent studies increasingly require integrative analysis across multiple samples, patients, or experimental conditions. In such settings, batch effects arising from technical variability between samples must be appropriately accounted for. NicheCompass incorporates a batch correction mechanism that enables joint embedding of cells from multiple samples into a shared latent space[10], and Nicheformer leverages a foundation model approach pretrained on large-scale single-cell and spatial atlases to achieve cross-dataset generalization[23]. Integrating similar multi-sample harmonization strategies into Mievformer would broaden its applicability to cohort-level studies and facilitate the identification of microenvironmental changes associated with disease progression or treatment response. Addressing these limitations represents promising directions for improving spatial omics analysis methods.

## 4 Methods

### 4.1 In situ gene expression profiling

In situ RNA expression analysis was performed using Xenium (10x Genomics, CG000601). FFPE sections (10 *µ*m thick) were prepared on Xenium slides. Fixation and tissue permeabilization were performed according to the protocol provided by 10x Genomics. The probe hybridization mixture was prepared using a Xenium Human Multi-Tissue and Cancer Panel. The prepared probes were hybridized, and post-hybridization washing, ligation, and amplification were performed according to the user guide. Autofluorescence quenching and nuclear staining were conducted in the dark. Using the prepared slides, fluorescent probe hybridization and imaging were conducted via a Xenium Analyzer (10x Genomics). After the Xenium run, H&E staining was performed on the Xenium slide.

### 4.2 Data preprocessing

In this study, we analyzed spatial transcriptome data generated by three platforms: Xenium (10x Genomics), Stereo-seq (BGI Genomics), and Visium HD (10x Genomics). Each dataset underwent the following preprocessing pipeline using rapids-singlcell [24], a GPU-accelerated implementation of scanpy[25]. First, quality control (QC) metrics were calculated for each cell, and cells were filtered based on the number of detected genes. Specifically, cells with fewer than 20 detected genes were excluded. Next, gene filtering was performed to remove genes detected in fewer than 0.5% of cells to eliminate uninformative genes. Expression values were then normalized using total-count normalization, where each cell’s counts were scaled so that the total count per cell equals the median total count across all cells, followed by natural logarithm transformation via log(*x* + 1) (`log1p`). PCA was applied to the preprocessed expression matrix, retaining the first 30 principal components. A *k*-nearest neighbor graph was constructed in the PCA space, followed by UMAP[26] dimensionality reduction for visualization and Leiden clustering[27] for cell-type annotation. These clustering results were used to define cell populations for downstream microenvironmental analyses. The annotation was performed based on known marker genes for each cell type, which were derived from PanglaoDB[28]. Spatial coordinates were normalized by the mean distance to *k*-nearest neighbors (default *k* = 100) used in Mievformer, ensuring that relative positional coordinates were within an appropriate range for model input.

### 4.3 Datasets

We used five spatial transcriptomics datasets in this study. The **Xenium Lung** and **Xenium Pancreas** datasets were generated in this study using the Xenium In Situ platform (see “In situ gene expression profiling” for experimental details). These data will be deposited in the Gene Expression Omnibus (GEO) database upon acceptance. The **Xenium GBM** (glioblastoma) data were obtained from the 10x Genomics public dataset repository (https://www.10xgenomics.com/datasets). The **Stereo-seq Mouse Brain** data were obtained from the MOSTA project[15] via the STOmicsDB database (https://db.cngb.org/stomics/mosta/). The **Visium HD Mouse Brain** data were obtained from the 10x Genomics public dataset repository (https://www.10xgenomics.com/datasets).

### 4.4 Mievformer

#### 4.4.1 Input data preparation

Mievformer input is comprised by spatial transcriptome data containing gene expression vectors *y*_*i*_ and spatial coordinates *x*_*i*_ for each cell *i*, and computes microenvironmental embeddings *e*_*i*_ representing the local microenvironmental state. For computational efficiency and optimization stability, rather than operating directly on gene expression, Mievformer uses low-dimensional representations *z*_*i*_ obtained via PCA.

For each cell *i*, the embedding *e*_*i*_ is computed from the cell states {*z*_*j*_ | *j* ∈ *N* (*i*)} within its spatial neighborhood *N* (*i*), where *N* (*i*) denotes the set of neighboring cells excluding *i*. The neighborhood *N* (*i*) is determined using scikit-learn’s[29] `NearestNeighbors` module, with a default of 100 neighboring cells. Spatial coordinates are encoded as relative positions *d*_*ij*_ = *x*_*j*_ − *x*_*i*_ and transformed into positional encodings *δ*_*ij*_ via sinusoidal encoding[30], defined as:

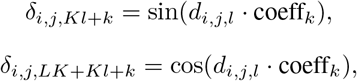

where coeff_*k*_ = exp(−2*k* log(10000)*/d*_model_), *d*_model_ denotes the embedding dimension, *L* is the dimensionality of spatial coordinates (*L* = 2 in this study), *K* = *d*_model_*/*(2*L*), and indices run over *k* = 0, 1, …, *K* − 1 and *l* = 0, …, *L* − 1.

### 4.4.2 Mievformer algorithm

Mievformer employs a masked self-supervised learning framework to compute the probability of observing a central cell state given its surrounding cellular context, and maximizes this probability during training. The architecture comprises NicheEncoder and Distributor as main components.

NicheEncoder constructs the input sequence by placing a learnable mask token at the central cell position: [mask + *δ*_*i,i*_, {*h*(*z*_*j*_) + *δ*_*ij*_ *j* | ∈ *N* (*i*)}]. Each neighboring cell state *z*_*j*_ is linearly projected to *d*_model_ dimensions (default: 64) via transformation *g* and combined with its positional encoding *δ*_*ij*_.

This sequence is processed through multiple Transformer encoder layers (default: 3), implemented using PyTorch’s[31] `nn.TransformerEncoderLayer`, which integrates multi-head self-attention and feedforward networks. The final hidden state of the mask token is transformed into a microenvironmental state *e*_*i*_ using a feedforward neural network.

Distributor computes the probability that each cell embedding *z*_*j*_ (*j* = 1, …, *M*) in the mini-batch could be observed at the central position given microenvironment *e*_*i*_. The model defines a score function:

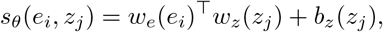

where *w*_*e*_ and *w*_*z*_ are neural networks mapping microenvironmental and cell-state embeddings to a shared hidden dimension *h*, and *b*_*z*_ provides a cell-state-dependent bias. We did not include bias term for a microenvironmental state *e*_*i*_, since the calculated scores are normalized for each *e*_*i*_. The conditional probability is obtained via softmax normalization:

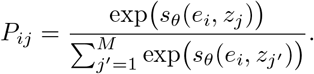

The training objective maximizes the likelihood that the observed central cell state *z*_*i*_ is generated from its inferred microenvironment *e*_*i*_ by minimizing the negative log-likelihood:

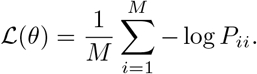

This formulation corresponds to the InfoNCE loss, a contrastive learning objective widely used in self-supervised representation learning[14]. InfoNCE maximizes mutual information between paired observations by treating one sample as positive and others in the batch as negatives. Hence, Mievformer’s training encourages learning microenvironmental embeddings that have high mutual information with their corresponding central cell states.

Model parameters are optimized using the `AdamW` optimizer[32] with a learning rate of 1*×*10^−3^, implemented via PyTorch Lightning[33]. Training is performed for a maximum of 1000 epochs with early stopping based on validation loss, using a patience of 20 epochs. The validation set is constructed by extracting a spatially contiguous central region from the tissue, specifically the central square region whose side length is 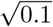 times the average of the tissue width and height, corresponding to approximately 10% of the total tissue area. This spatial validation strategy ensures that the model is evaluated on cells with spatial context distinct from the training set while maintaining biological coherence. Mini-batch size is set to 512 cells by default. The model checkpoint with the lowest validation loss is retained for downstream analysis. All neural networks were implemented in PyTorch[31].

#### 4.4.3 Relationship between score function and density ratio obtained by Mievformer

In this section, we demonstrate that the score function *s*_*θ*_(*e, z*) learned by Mievformer in the previous section approximates the conditional density ratio *p*(*z* | *e*)*/p*(*z*), which quantifies the degree to which the observing cell state *z* becomes more probable when it is conditioned on the microenvironment *e*, and represents the mutual information between microenvironment *e* and cell state *z*.

Starting from the score function *s*_*θ*_(*e, z*) defined in the previous section, we construct the positive ratio function and its normalization:

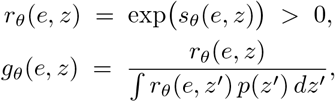

where *p*(*z*) denotes the true marginal distribution over cell embeddings. By construction, *g*_*θ*_(*e, z*) satisfies the normalization constraint

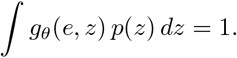

Consequently, we can define a valid probability distribution over *z* conditioned on *e* as:

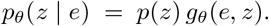

We now show that minimizing the Kullback-Leibler divergence between the true conditional distribution *p*(*z* | *e*) and the model distribution *p*_*θ*_(*z* | *e*) is equivalent to minimizing the training loss defined in the previous section. The expected KL divergence can be expanded as:

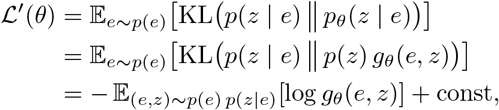

where the constant term corresponds to the entropy of *p*(*z* | *e*) and is independent of *θ*.

Here, we approximate this expected loss using a mini-batch of *M* observed cell-microenvironment pairs 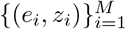:

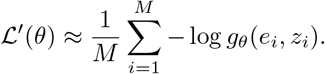

Additionally, the normalization integral in *g*_*θ*_(*e, z*) is approximated by Monte Carlo sampling over the mini-batch:

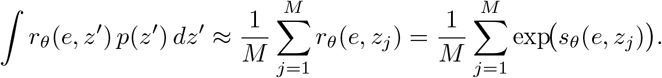

Substituting this approximation yields

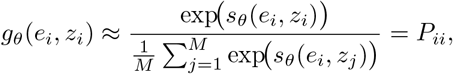

where *P*_*ii*_ is the softmax probability defined in the previous section. Therefore, the expanded KL divergence results in

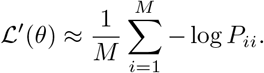

which is identical to the loss function presented in the Mievformer algorithm section.

This equivalence demonstrates that minimizing the training loss of Mievformer is mathematically equivalent to minimizing the KL divergence between *p*(*z* | *e*) and *p*_*θ*_(*z* | *e*). At the optimal solution, the model exactly recovers the true conditional:

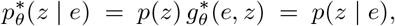

And, therefore,

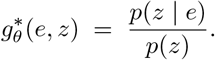

#### 4.4.4 Estimation of cell population density

The cell population density *ρ*_*C*_(*e*) in a given microenvironment *e* can be derived by integrating the learned conditional distribution *p*_*θ*_(*z* | *e*) across all cell states belonging to population *C*, utilizing the density ratio relationship established in the previous section.

The density of population *C* in microenvironment *e* is defined as:

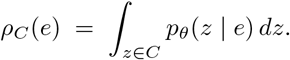

Substituting the density ratio decomposition *p*_*θ*_(*z* | *e*) = *p*(*z*) *g*_*θ*_(*e, z*) from the previous section, we obtain

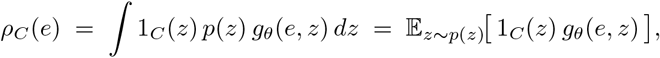

where 1_*C*_(*z*) denotes the indicator function for membership in population *C*.

In practice, we approximate this expectation using the empirical distribution 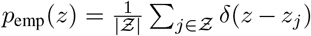, where *Ƶ* denotes the set of all observed cell embeddings. This yields

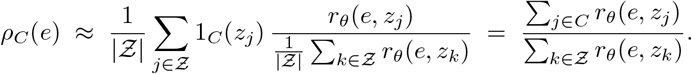

Thus, under the empirical distribution, the density is the ratio of the sum of *r*_*θ*_(*e, z*) = exp(*s*_*θ*_(*e, z*)) over population *C* to the sum over the entire dataset *Ƶ*.

For computational efficiency with large-scale datasets, we employ Monte Carlo approximation via independent random sampling from *C* and *Ƶ*. Let *U*_*C*_ ⊂ *C* and *U*_*ƶ*_ ⊂ *Ƶ* denote simple random samples of sizes *M*_*C*_ and *M*_*ƶ*_, respectively. The full sums can be approximated by unbiased Monte Carlo estimators:

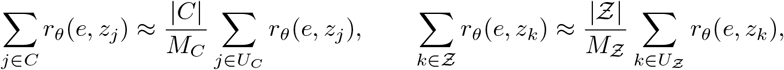

yielding the density estimate

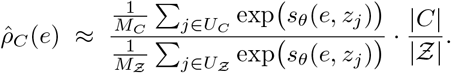

By construction, *ρ*_*C*_(*e*) ∈ [0, 1].

For numerical stability, we compute the estimate in log-space using the log-sum-exp (LSE) function:

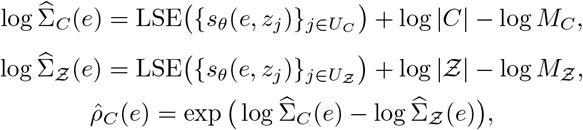

where LSE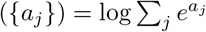.

#### 4.4.5 Single-cell level microenvironment distribution estimation

For each cell embedding *z*, we estimate the posterior distribution across microenvironments through density-ratio-based Bayesian inversion. We define the joint distribution of cell embedding *z* and microenvironment *e* as:

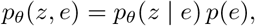

where *p*(*e*) denotes the true distribution of microenvironments. By Bayes’ theorem,

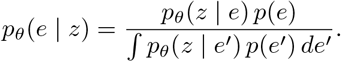

Introducing the density ratio function 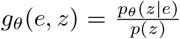, which is the function learned by Mievformer as established in the previous section, we obtain:

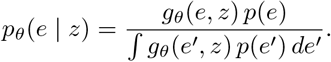

Therefore, the density-ratio-based Bayesian inversion yields:

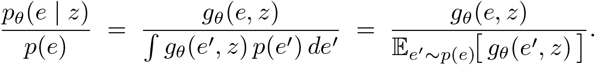

In practice, we approximate this expectation using the empirical distribution 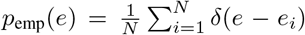 where 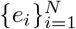 are the observed microenvironmental embeddings:

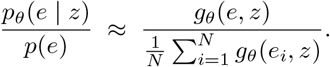

This expression demonstrates that bidirectional probabilistic inference (*e ↔ z*) is unified through the density ratio framework: if *g*_*θ*_(*e, z*) approximates *p*_*θ*_(*z* | *e*)*/p*(*z*), it can be directly used for estimating *p*_*θ*_(*e* | *z*)*/p*(*e*).

To obtain the posterior probability for a microenvironmental cluster *c*, we define 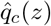 as the integral of *p*_*θ*_(*e*|*z*) across the region where *e* belongs to cluster *c*:

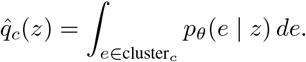

Using the density ratio relationship derived above, we can rewrite this as:

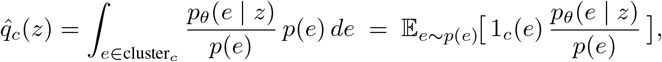

where 1_*c*_(*e*) = 1 if *e* ∈ cluster_*c*_ and 0.

In practice, we approximate this expectation using the empirical distribution 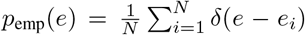, where 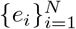 are the observed microenvironmental embeddings:

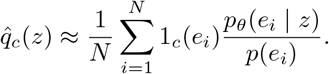

Substituting the previously derived density ratio approximation 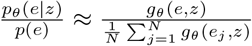, we obtain:

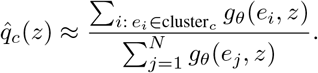

This expression directly yields the posterior probability of cluster membership, as 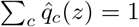 by construction. For computational efficiency with large datasets, we can further approximate this using Monte Carlo sampling: drawing a random subset *U*_*c*_ ⊂ {*i* : *e*_*i*_ ∈ cluster_*c*_} of size *M*_*c*_ and a random subset *U*_*ƶ*_ ⊂ {1, …, *N*} of size *M*_*Z*_, we obtain:

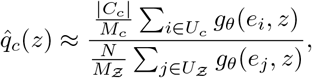

where |*C*_*c*_| denotes the number of microenvironments in cluster *c*.

This density-ratio-based framework enables rigorous estimation of each cell’s microenvironmental distribution profile, facilitating cell clustering according to microenvironmental preferences and quantitative characterization of context-dependent cellular behavior.

### 4.5 Evaluation metrics

For quantitative evaluation of microenvironmental representations, the following metrics were employed:

#### Niche Adjusted Rand Index (Niche ARI)

Evaluates the concordance between ground-truth clusters and estimated microenvironmental clusters in simulation data. ARI ranges from 0 to 1, with higher values indicating greater agreement. For clustering microenvironmental representations, *k*-means was applied with the number of clusters set to match the true number of microenvironmental clusters configured in the simulation.

#### Cell Type Affinity Similarity (CAS)

Assesses the concordance between cell-type colocalization patterns in microenvironmental embedding neighborhoods and those in spatial neighborhoods. Specifically, *k*-nearest neigh-bor graphs (default *k* = 5) were constructed for both microenvironmental embeddings and spatial coordinates using rapids_singlecell[24], and neighborhood enrichment matrices were computed using squidpy’s[34] `nhood_enrichment` function. The resulting matrices were flattened, and Spearman’s rank correlation coefficient was calculated using scipy.stats’[35] `spearmanr` function.

#### Shuffled Cell Type Affinity Similarity (SCAS)

Evaluates the preservation of colocalization patterns when spatial coordinates are randomly permuted within microenvironmental clusters. *K*-means clustering was applied to microenvironmental embeddings (default *K* = 50), and spatial coordinates were shuffled within each cluster. Neighborhood enrichment matrices were computed for both original and shuffled spatial coordinates, and the mean Pearson correlation coefficient was calculated row-wise across cell types.

#### Neighbourhood Enrichment Accuracy (NEA)

Defined as the Pearson correlation coefficient between the following two matrices: (1) Spatial enrichment matrix *E*_sp_ ∈ ℝ^*C×C*^ : Let *j* = NN_sp_(*i*) denote the spatial nearest neighbor of cell *i*, and let *y*_*i*_ ∈ {1, …, *C*} denote the cell type. Define *n*_*c*_ = # {*i* | *y*_*i*_ = *c*} as the number of cells of type *c*, and the co-occurrence count of type pairs as

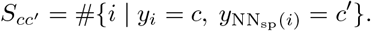

Then,

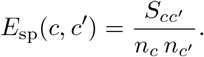

(2) Expected enrichment matrix based on microenvironmental clusters *E*_niche_ ∈ ℝ^*C×C*^ : *K*-means clustering (default *K* = 50) was applied to the latent representation to obtain microenvironmental cluster assignments *t*_*i*_ ∈ {1, …, *K*}. From the type-by-cluster frequency matrix

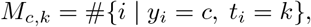

we constructed the cluster composition matrix for each cell type

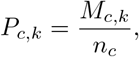

and defined

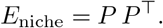

The final score is computed as

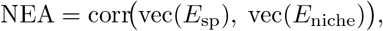

where corr and vec denote Pearson correlation and vectorization, respectively.

#### Dual Recovery (DREC)

Quantifies the proportion of cells for which expression-similar and spatially proximal neighbors belong to the same microenvironmental cluster. Specifically, for each cell, the 10th nearest neighbor cell was selected based on expression differences. DREC was computed as the fraction of cases where the spatial nearest neighbor of that cell belongs to the same microenvironmental cluster as the original cell. *K*-means clustering was used for microenvironmental clustering with the number of clusters set to 50 by default.

Among these metrics, DREC, which exhibited the highest correlation with ground-truth evaluation (Niche ARI) in simulation data, was adopted as the primary performance evaluation metric for real datasets. For unsupervised evaluation, a relatively large number of microenvironmental clusters (default *K* = 50) was used to reduce performance variability due to the clustering results.

In addition to the metrics described above, we evaluated four metrics from the NicheCompass benchmarking module[10]: Graph Connectivity Similarity (NCB-GCS), Cell-type Affinity Similarity (NCB-CAS), Niche Average Silhouette Width (NCB-NASW), and Maximum Leiden Adjusted Mutual Information (NCB-MLAMI). For detailed definitions of these metrics, see Birk et al.[10].

### 4.6 Comparison methods

#### NicheCompass[10]

A microenvironmental representation learning method based on graph attention networks (GAT). The method integrates information on cell-cell interaction and other biological processes using gene program dictionaries obtained from OmniPath[36], MEBOCOST[37], and NicheNet[38], constructing a spatial graph with four neighbors per cell. Learning was performed using a `GATv2Conv`[11] encoder with learning rate 0.001, a maximum of 400 epochs, and early stopping. Microenvironmental representations were obtained by setting the active gene program threshold ratio to 0.01.

#### Banksy[12]

A microenvironmental representation method that integrates mean expression profiles and spatial expression gradients within local neighborhoods. Scale-Gaussian weights were applied with 15 neighbors, and the spatial regularization parameter *λ* was set to 0.8. PCA was applied to the resulting features to obtain a 20-dimensional representation.

#### STAGATE[8]

A spatial transcriptomics analysis method based on graph convolutional neural networks. Spatial binning on a 50*×*50 grid was applied to reduce memory requirements. The spatial network radius cutoff was set to 400, and pure representation learning was executed with the autoencoder weight parameter *α* = 0.

#### GraphST[9]

A graph neural network-based representation learning method for spatial transcriptomics data. Spatial binning on a 100*×*100 grid was applied to reduce memory requirements. Latent representations are learned through graph convolution utilizing spatial proximity-based graph structures.

#### COVET[39]

Spatial omics features derived from ENVI, an integrated analysis framework for spatial and single-cell omics based on variational autoencoders[40]. Spatial embeddings were computed using the `scenvi.utils.compute_covet()` function, followed by PCA-based dimensionality reduction to 20 dimensions to obtain the final microenvironmental representation.

#### CellCharter[13]

A spatial cell niche identification method that aggregates neighborhood features across multiple graph layers. A spatial neighbor graph was constructed using Delaunay triangulation via Squidpy[34]. PCA was applied to gene expression data to obtain 50-dimensional representations, which were then aggregated over the spatial graph using `cc.gr.aggregate_neighbors()` with `n_layers=3` to capture multi-scale neighborhood information. The resulting aggregated features were used as microenvironmental representations.

As comparison baselines, the following methods were also included:

#### PCA

Principal component analysis was applied to gene expression data to obtain a 20-dimensional representation. This serves as a basic baseline using expression information alone.

#### Spatial coordinates

Raw spatial coordinates were used directly as microenvironmental representations. This baseline evaluates the performance limits of representations that rely solely on spatial information.

### 4.7 Simulation data generation with multiple microenvironmental patterns

To enable quantitative evaluation and benchmarking of Mievformer against competing methods, we generated synthetic spatial transcriptomics data with known ground-truth microenvironmental structure. The simulation framework supports multiple distinct microenvironmental patterns while preserving realistic gene expression characteristics by directly sampling expression profiles from observed cells in pancreatic cancer Xenium data.

#### 4.7.1 Spatial structure and microenvironmental pattern definition

The simulation domain was defined as a square region of dimensions 1000 × 1000 arbitrary units. We configured *D* distinct microenvironmental patterns (*D* = `duct_category_num`), each representing a spatially contiguous region with defined cellular composition.

For each pattern *k* ∈ {0, …, *D* − 1}, we generated *n*_duct_ annular structures (ducts) as follows:

- Duct centers were sampled uniformly within the simulation domain, and each was randomly assigned a pattern label *k* uniformly.
- To introduce spatial heterogeneity, the radius *r* and boundary thickness *t* of each duct were perturbed from baseline values (`radius_base` and `thick_base`, respectively) according to:

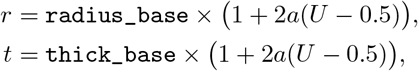

where *U ∼* Uniform[0, 1] and *a* = 0.2 controls the magnitude of variation. The inner and outer radii were computed as *r*_in_ = max(0, *r* − *t*) and *r*_out_ = *r* + *t*, defining three concentric microenvironmental zones per duct: inner, boundary, and outer.

Each spatial location was assigned one of 2*D* + 1 ground-truth microenvironmental labels: outside (background), inner_*k*, or boundary_*k* for *k* = 0, …, *D* − 1.

### 4.7.2 Cell placement and microenvironmental assignment

A total of *n*_cell_ cells were uniformly distributed across the simulation domain. Each cell’s microenvironmental label was determined via nearest-neighbor search as follows:

- For each pattern *k*, we constructed a *k*-nearest neighbors index (*k* = 10) using only duct centers of that pattern and computed the distance from each cell to its nearest duct of pattern *k*.
- If the minimum distance was less than *r*_out_, the cell was provisionally labeled as boundary_*k*; if less than *r*_in_, it was relabeled as inner_*k*.
- Labels were assigned in hierarchical order (boundary *→* inner) across all patterns, with later patterns potentially overwriting earlier assignments.
- Cells whose distances to all ducts exceeded *r*_out_ retained the default label Outside.

#### 4.7.3 Cell-type composition and expression profile sampling

For each microenvironmental label (outside, inner_*k*, and boundary_*k*), we defined distinct cell-type compositions and sampled corresponding expression profiles from the reference dataset.

- We first computed the empirical cell-type frequency vector *π* from the reference Xenium data using a predefined cell-type annotation key (`cluster_key`).
- For each microenvironment *n*, a cell-type composition vector *p*^(*n*)^ was sampled from a Dirichlet distribution with concentration parameter *α* = `alpha_base` × *π*, where `alpha_base` controls the degree of variation around the reference composition.
- For each cell assigned to microenvironment *n*, a cell type was drawn according to the multinomial distribution Multinomial(1, *p*^(*n*)^).
- The gene expression profile for that cell was obtained by randomly selecting one observed cell of the assigned type from the reference dataset and copying its preprocessed expression vector.

This sampling strategy ensures that simulated expression profiles reflect the biological variability present in real data while maintaining controlled microenvironmental structure.

#### 4.7.4 Parameter settings

Unless otherwise specified, simulations employed the following default parameters: `alpha_base` = 5, *n*_duct_ = 1000, *n*_cell_ = 300,000, `radius_base` = 10, `thick_base` = 2, and *D* = `duct_category_num` (varied across experiments). The relative perturbation magnitude was fixed at *a* = 0.2, and *k* = 10 nearest neighbors were used for microenvironmental assignments. These parameters were externally configurable via model identifiers in the simulation script (`make_duct_sim.py`).

### 4.8 Immunohistochemistry

Paraffin-embedded tissue sections (4 µm) were deparaffinized, rehydrated, and subjected to antigen retrieval using Target Retrieval Solution, Citrate pH 6 (Dako). Endogenous peroxidase activity was blocked with 3% hydrogen peroxide in methanol for 5 min, followed by blocking with 3% BSA in PBS for 30 min. Sections were incubated overnight at 4 °C with an anti–c-Myc antibody (Y59, Abcam), then with EnVision+ Dual Link System-HRP (Dako) for 30 min at room temperature. Signals were visualized using the Liquid DAB+ Substrate Chromogen System (Dako), and sections were counterstained with Mayer’s hematoxylin, dehydrated, cleared, and mounted. Hematoxylin and eosin staining was performed for background evaluation.

Quantitative analysis of c-Myc immunohistochemistry was performed to compare expression levels between fibroblast-proximal and tumor-core regions. Color deconvolution was applied to separate hematoxylin and DAB channels using the HED color space. Nuclear segmentation was performed on the hematoxylin channel using Otsu’s thresholding (multiplier = 0.6) followed by the watershed algorithm. The DAB positivity threshold was determined by fitting a two-component Gaussian Mixture Model to pixel values from all nuclear regions, with the threshold set at the midpoint between component means. For each nucleus, the DAB-positive pixel fraction was calculated as the proportion of pixels exceeding this threshold. Two regions of interest (130 × 130 pixels) were defined in fibroblast-proximal and tumor-core regions. Statistical comparison was performed using the Mann–Whitney *U* test.

## Acknowledgements

This work was supported by P-PROMOTE (grant no. 24ama221609h000 to Y.K.) and PRIME (grant no. 25gm7010014h0001 to Y.K.) from the Japan Agency for Medical Research and Development (AMED), the Moonshot Research and Development Program (grant no. JP22zf0127009 to H.M.), National Cancer Center Research and Development Fund (grant no. 2024-A-6 to Y.K.), and Grant-in-Aid for Scientific Research (grant no. 23K16991 to Y.K.) from the Japan Society for the Promotion of Science (JSPS).

## Author contributions

Y.K. designed the study and performed software development and data analysis. Y.T., F.C. and H.M. conducted and supervised experiments. H.H., S.H., M.I., and T.S. performed software development and data analysis. H.H. and K.I. performed data analysis.

## Data and code availability

The Xenium Lung and Xenium Pancreas datasets generated in this study will be deposited in the Gene Expression Omnibus (GEO) database upon acceptance. The Stereo-seq Mouse Brain data are available from the STOmicsDB (https://db.cngb.org/stomics/mosta/). The Xenium GBM and Visium HD Mouse Brain datasets are available from the 10x Genomics public dataset repository (https://www.10xgenomics.com/datasets). The source code for Mievformer is publicly available as a Python package at https://github.com/kojikoji/mievformer_package, with documentation at https://kojikoji.github.io/mievformer_package/index.html.

