## Supplementary figures for "Probabilistic coupling of cellular and microenvironmental heterogeneity by masked self-supervised learning"

### Supplementary Materials for Probabilistic coupling of cellular and microenvironmental heterogeneity by masked self-supervised learning

Yasuhiro Kojima[1,†,*], Yosuke Tanaka[2,†], Haruka Hirose[1,3], Fumiko Chiwaki[4], Kazuya Nishimura[1,5], Shuto Hayashi[3,6], Kota Itahashi[7], Masato Ishikawa[8], Teppei Shimamura[3], Hiroyuki Mano[9]

[1]Laboratory of Computational Life Science, National Cancer Center Research Institute, Tokyo, Japan
[2]Laboratory of Cancer Target Discovery, National Cancer Center Research Institute, Tokyo, Japan
[3]Department of Computational and Systems Biology, Institute of Science Tokyo, Bunkyo-ku, Tokyo, Japan
[4]Department of Translational Oncology, National Cancer Center Research Institute, Tokyo, Japan
[5]D3 center, The University of Osaka, Suita, Osaka, Japan
[6]Innovation Center of NanoMedicine, Kawasaki Institute of Industrial Promotion, Kawasaki, Kanagawa, Japan
[7]Division of Cancer Immunology, Research Institute/Exploratory Oncology Research & Clinical Trial Center (EPOC), National Cancer Center, Tokyo, Japan
[8]Institute for Life and Medical Sciences, Kyoto University, Kyoto, Japan
[9]Division of Cellular Signaling, National Cancer Center Research Institute, Tokyo, Japan

[†]These authors contributed equally to this work.
[*]Corresponding author: Yasuhiro Kojima

#### Supplementary Figures

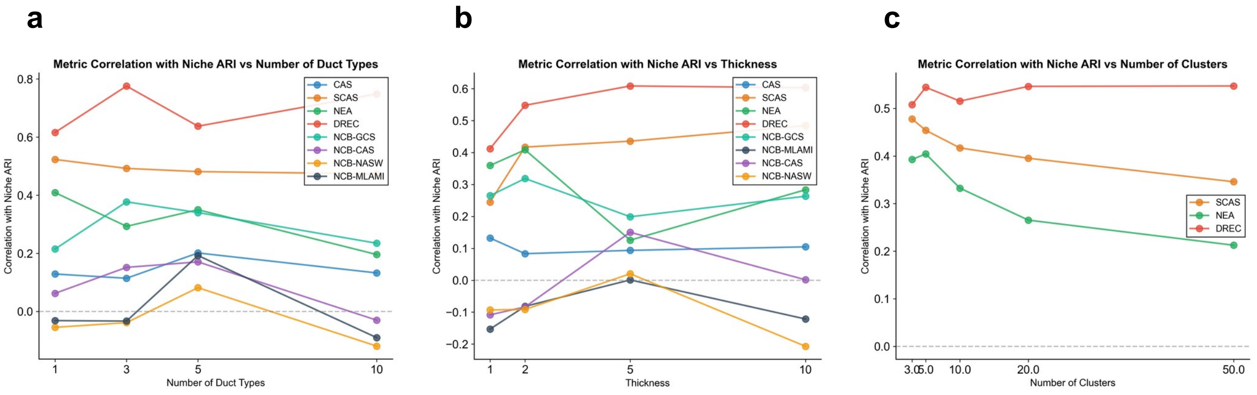


Figure 1: **Metric Correlation Under Varying Simulation Conditions.** **(a)** Line plot showing Pearson correlation between eight evaluation metrics (CAS, SCAS, NEA, DREC, NCB-GCS, NCB-CAS, NCB-NASW, NCB-MLAMI) and ground-truth Niche ARI as a function of number of duct types (dcat). **(b)** Line plot showing Pearson correlation between eight evaluation metrics and ground-truth Niche ARI as a function of duct wall thickness. **(c)** Line plot showing Pearson correlation between three metrics (SCAS, NEA, DREC) and ground-truth Niche ARI as a function of number of clusters.

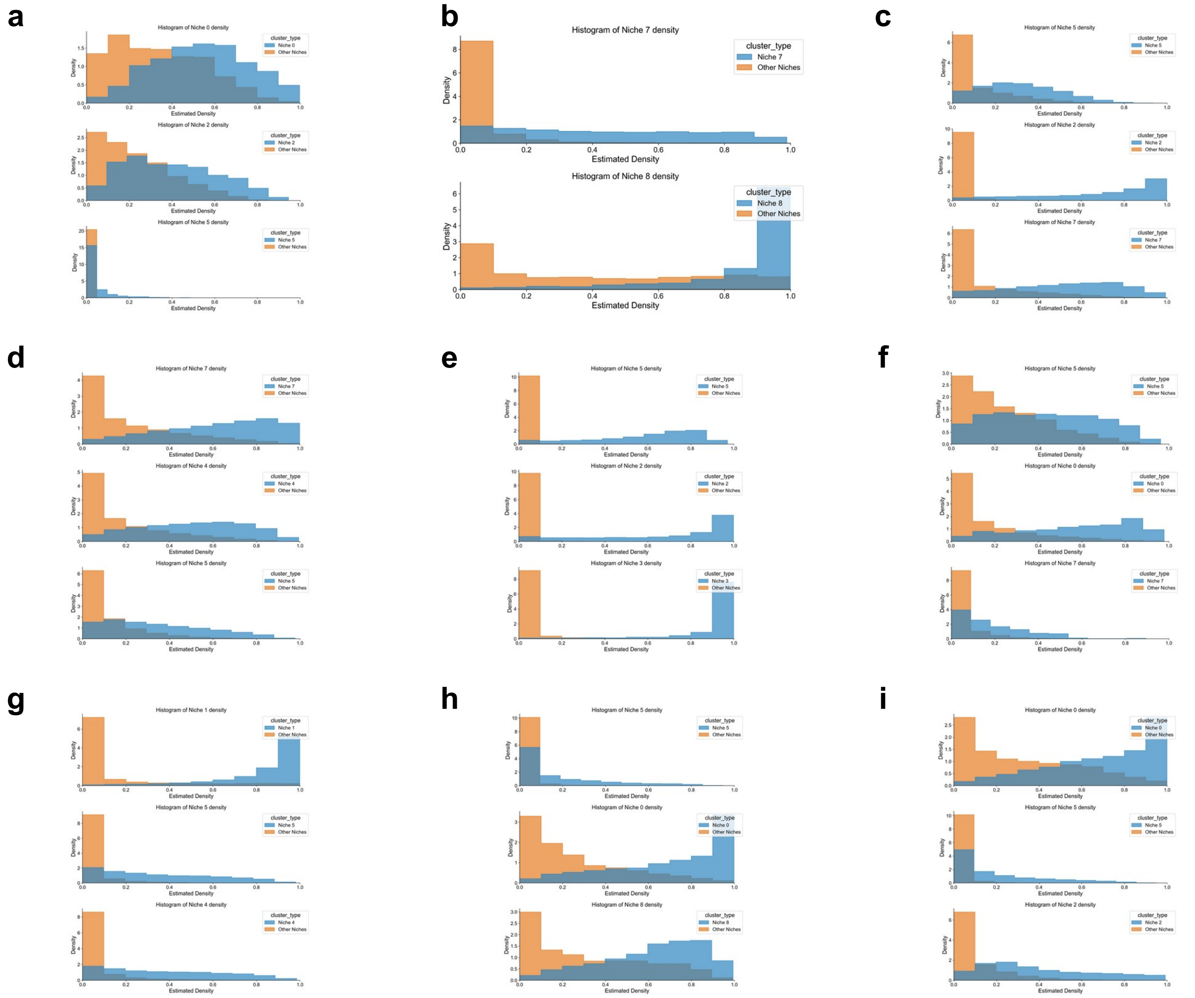


Figure 2: **Density Aggregation Histograms for All Cell Types.** Each panel shows a histogram comparing estimated population density distributions between the top 3 assigned niche clusters (blue) and other niche clusters (orange) for the following cell types: **(a)** B cells, **(b)** Ciliated Epithelial cells, **(c)** Endothelial cells and Pericytes, **(d)** Epithelial cells, **(e)** Fibroblasts and Smooth Muscle cells, **(f)** Mast cells, **(g)** Monocytes and Macrophages, **(h)** Plasma cells, **(i)** T cells.

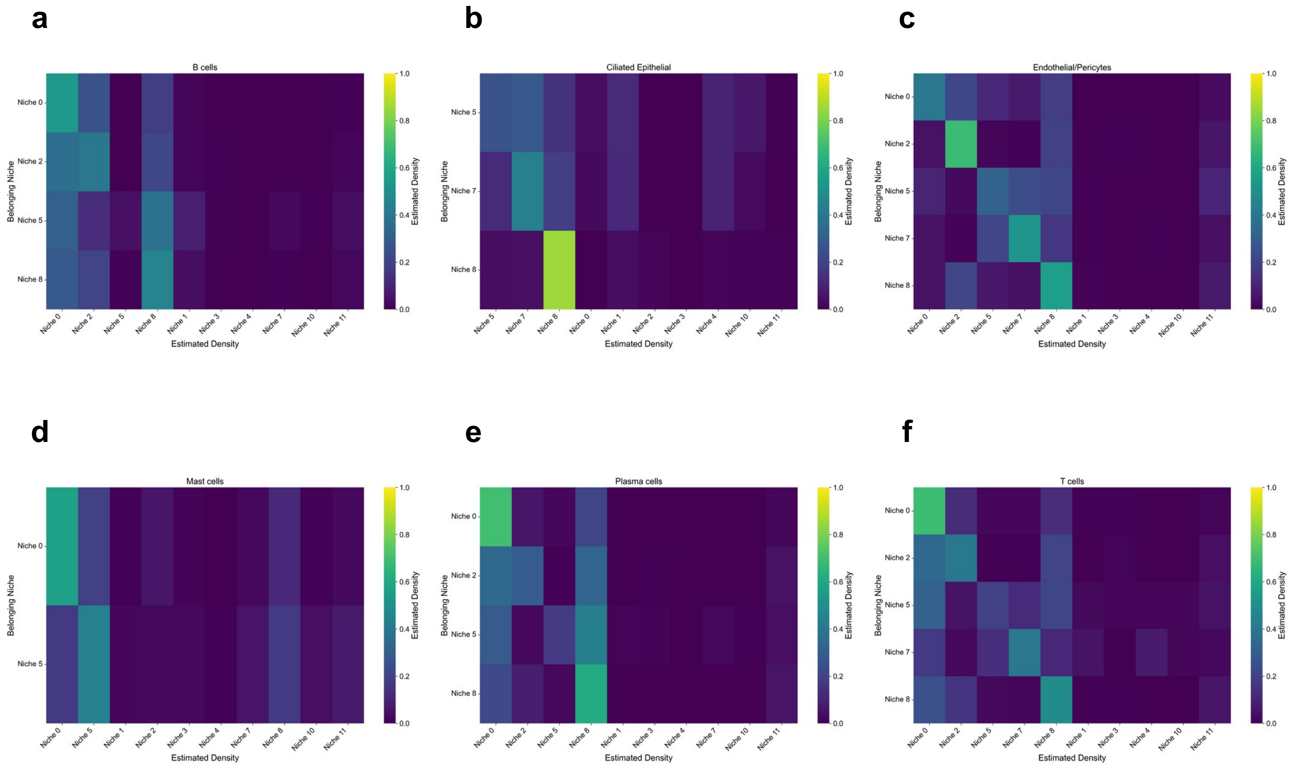


Figure 3: **Density Aggregation Heatmaps for Additional Cell Types.** Each panel shows a hierarchical clustering heatmap of estimated posterior distribution across niche clusters. Rows represent belonging niche clusters; columns represent estimated density for each niche cluster. Cell types shown: **(a)** B cells, **(b)** Ciliated Epithelial cells, **(c)** Endothelial cells and Pericytes, **(d)** Mast cells, **(e)** Plasma cells, **(f)** T cells.

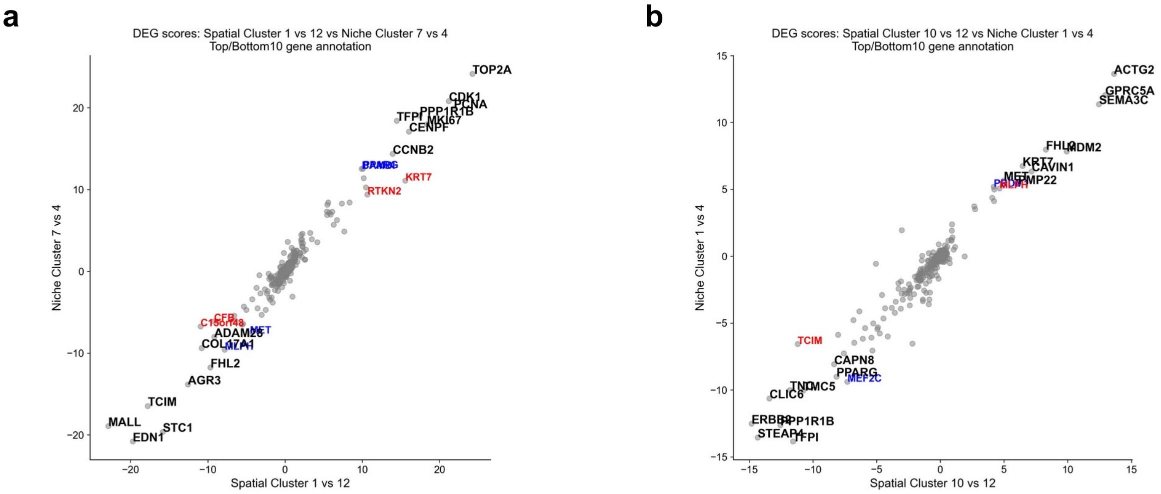


Figure 4: **Differentially Expressed Gene Scores Across Niche Clusters.** Each panel shows a scatter plot of differentially expressed gene scores. X-axis: DEG scores between spatial clusters. Y-axis: DEG scores between niche clusters. Top and bottom 10 genes are annotated; red indicates genes upregulated in both comparisons, blue indicates genes downregulated. **(a)** Dist 1 vs 12 and Niche 7 vs 4. **(b)** Dist 10 vs 12 and Niche 1 vs 4.

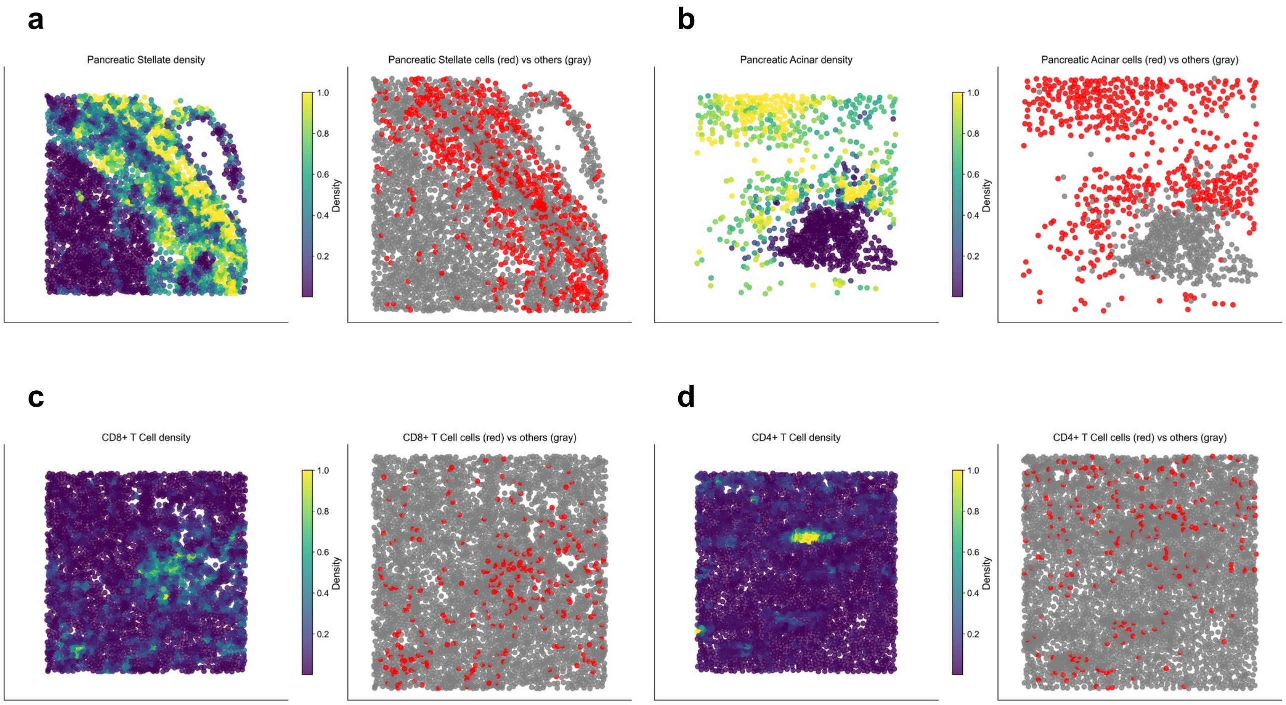


Figure 5: **Spatial Distribution of Additional Cell Types in Pancreatic Tissue.** Each panel shows the spatial distribution of a cell type. Left: estimated population density for Mievformer's latent representation at each single cell spot. Right: cell positions (red=target cell type, gray=others). Cell types shown: **(a)** pancreatic stellate cells, **(b)** pancreatic acinar cells, **(c)** CD8-positive alpha-beta T cells, **(d)** CD4-positive alpha-beta T cells.
